# Temperature Evolution Following Joint Loading Promotes Chondrogenesis by Synergistic Cues via Calcium Signaling

**DOI:** 10.1101/2021.06.29.450339

**Authors:** Naser Nasrollahzadeh, Peyman Karami, Jian Wang, Lida Bagheri, Yanheng Guo, Philippe Abdel-Sayed, Lee Ann Applegate, Dominique P. Pioletti

**Affiliations:** Laboratory of Biomechanical Orthopedics, Institute of Bioengineering, EPFL, Switzerland; Institut des Matériaux et Institut des Sciences et Ingénierie Chimiques, Laboratoire des Polymères, EPFL, Bâtiment MXD, Station 12, CH-1015 Lausanne, Switzerland; Regenerative Therapy Unit, Department of Musculoskeletal Medicine, Lausanne University Hospital (CHUV), Switzerland

**Keywords:** mechanobiology, tissue viscoelasticity, Cartilage self-heating, thermo-mechanotransduction, TRPV4, calcium signaling

## Abstract

During loading of viscoelastic tissues, part of the mechanical energy is transformed into heat that can locally increase the tissue temperature, a phenomenon known as self-heating. In the framework of mechanobiology, it has been accepted that cells react and adapt to mechanical stimuli. However, the cellular effect of temperature increase as a by-product of loading has been widely neglected. In this work, we focused on cartilage self-heating to present a “thermo-mechanobiological” paradigm, and demonstrate how the synergy of a biomimetic temperature evolution and mechanical loading could influence cell behavior. We thereby developed a customized *in vitro* system allowing to recapitulate pertinent *in vivo* physical cues and determined the cells chondrogenic response to thermal and/or mechanical stimuli. Cellular mechanisms of action and potential signaling pathways of thermo-mechanotransduction process were also investigated. We found that co-existence of thermo-mechanical cues had a superior effect on chondrogenic gene expression compared to either signal alone. Specifically, a synergetic effect was observed for upregulation of Sox9 by application of the physiological thermo-mechanical stimulus. Multimodal TRPV4 channels were identified as key mediators of thermo-mechanotransduction process, which becomes ineffective without external calcium sources. We also observed that the isolated temperature evolution, as a by-product of loading, is a contributing factor to the cells response and this could be considered as important as the conventional mechanical loading. Providing an optimal thermo-mechanical environment by synergy of heat and loading portrays new opportunity for development of novel treatments for cartilage regeneration and can furthermore signal key elements for emerging cell-based therapies.

## 1 Introduction

Mechanoregulation is a central process to maintain health and functionality of musculoskeletal tissues. Cells within the extracellular matrix (ECM) of load-bearing tissues receive various physical stimuli and maintain tissue homeostasis and function through reciprocal responses [1-3]. The externally applied load generates different sensible cues within the cells surrounding micro-environments, such as spatiotemporal deformation, interstitial fluid pressure, shear stress and ion mobility [2, 4]. In this process, mechanical properties, organizational structure and constituents of the ECM are massively influential and mediate the conveyed signals. Cell sensory systems integrate and convert the perceived physical inputs into the biochemical transients via different mechanisms of actions (e.g. calcium signaling [5]). Accordingly, cells react and the elicited response modifies expression of genes and subsequent proteins to adapt tissue composition and structure [1, 6]. In particular, chondrogenic cells were shown to respond to physical stimuli when various dynamic loading regimens were applied (compression, torsion, tension, hydrostatic pressure) [7, 8] and when scaffold materials with different properties were used (elastic, viscoelastic, porous, non-porous) [9, 10]. However, the role of multimodal physical cues and associated coupling phenomena in chondrogenesis still remains to be fully understood.

Articular cartilage is a load-bearing tissue with significant dissipative capacity. This property arises from the cartilage ECM intrinsic viscoelasticity and fluid-solid interactions [11, 12]. Dissipation is a function of the applied loading as well as material characteristics and is therefore an integrative variable for macroscopic physical cues generating spatiotemporal signals around cells. We have capitalized on this concept and demonstrated that matching dissipative capacity of scaffolds with native cartilage is an effective strategy to convey chondro-inductive cues to cells seeded in scaffolds under cyclic loading [13, 14]. Apart from direct effects of mechanical dissipation, being associated with matrix deformation, shear stress and interstitial fluid pressure around cells, the cartilage dissipation may indirectly influence chondrocyte behavior by increasing tissue temperature. This phenomenon is called self-heating and occurs following conversion of the lost mechanical energy to heat over time [15]. In addition to cartilage dissipation, other factors such as ambient temperature and heat transfer from surrounding tissues could contribute to the temperature evolution in the knee joint. Previous studies indicated that cyclic compression exerted on cartilage could significantly increase local temperature [15-17]. Notably, *in vivo* measurements inside intra-articular regions of knee joints have shown a temperature rise from 32°C (at rest) to ∼ 38°C following one hour of jogging activity [16] at the ambient temperature of 19°C. Chondrocytes are sensitive to temperature and the literature suggests that their biophysical response to heat stimulation is dose-dependent [17-19]. However, none of the conducted studies are based on a biomimetic temperature evolution regime. The loading induced self-heating of cartilage, as shown in Figure 1, is an unexplored coupled phenomenon in the field of biomechanics which could initiate a new “thermo-mechanobiology” paradigm.

**Figure 1.**
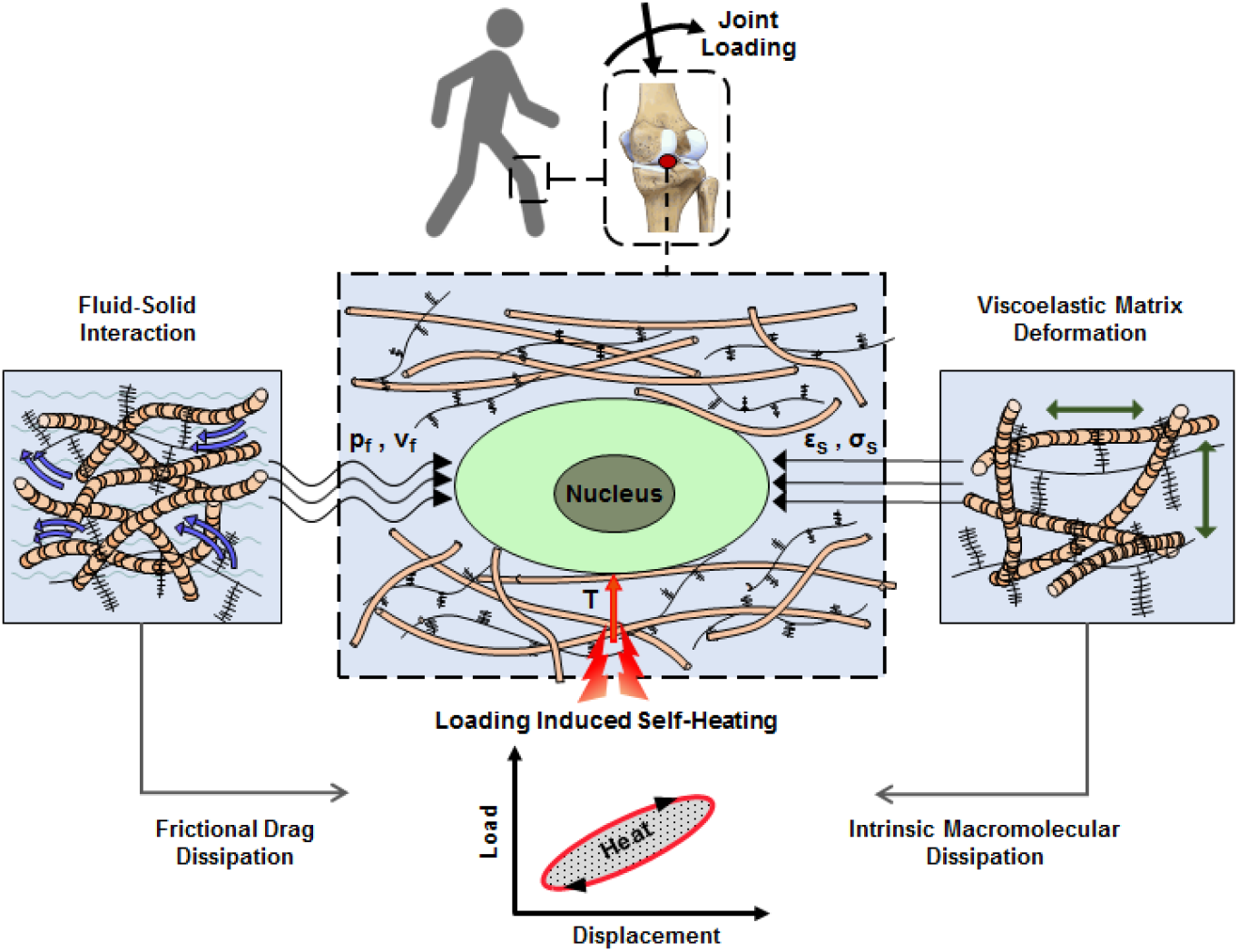
Cartilage loading induced self-heating during physical activity. Mechanical hysteresis in cartilage tissue during joint loading generates heat and causes the temperature rise over time. The self-heating of cartilage originates from both, intrinsic matrix viscoelasticity and solid-fluid interaction sources during joint loading. Structural and material characteristics of matrix microenvironment are acting in concert with external stresses to provide direct and indirect biophysical cues for cells such as deformation and temperature variation (p_f_, v_f_: fluid pressure and flow velocity; *ε*_*s*_, *σ*_*s*_ solid matrix strain and stress; T: temperature).

In cartilage microenvironment, different elements residing inside and/or on plasma membrane of chondrocytes (cytoskeleton, nucleus, ion channels, primary cilium, integrin and TGFβ_3_ receptors) could be activated by receiving spatiotemporal cues and trigger intracellular changes [7, 9, 20]. In particular, transient variation of intracellular calcium is a fundamental pathway for transduction of an external physical stimulus to control cellular functions [21]. It has been shown that, fluid flow [22], mechanical strain [23, 24], temperature [25], and osmolarity [5] modulate cytosolic calcium oscillation in chondrocytes. Among different calcium mediators, transient receptor potential vanilloid 4 (TRPV4) is a multimodal ion channel that significantly contributes to the transduction of physical cues [26-28]. The literature suggests that over-activation and full inhibition of TRPV4 channels in chondrocytes cause negative effects on joint development and cartilage homeostasis [29-32]. Of note, TRPV4 channels are responsive to mechanical loading [5, 24] and temperature rise [33, 34], making it a potential sensory system for translation of stimuli involved in loading induced self-heating phenomenon.

The overall objective of the present work is to address cartilage thermo-mechanobiology. For this purpose, we first developed an original model system capable of simulating the self-heating phenomenon *in vitro*. Specifically, our customized system encompassed a modular bioreactor designed to independently control the evolution of temperature and applied mechanical loading on cell-laden constructs, as well as a biomechanically functional scaffold recapitulating cartilage main properties. Having developed the tailored *in vitro* platform, we then determined the individual contribution and the combined effects of temperature and loading on cell response. In this study, the cellular reaction was evaluated through varied expression of chondrogenic genes following an applied stimulus. Finally, we aimed to identify downstream signaling and the potential role of TRPV4 ion channels as a mediator in the thermo-mechanotransduction process. Accordingly, the influence of thermo-mechanical cues on cell response was investigated when inhibiting TRPV4 channels as a signal transducer or diminishing cytosolic calcium variation as a mechanism of action. An insight on the effect of cartilage microenvironment on chondrocyte behavior and respective transduction cascade can positively contribute to the development of emerging cell-based therapies for tissue regeneration.

## 2 Results

### 2.1 A Custom-made *in vitro* Platform for Cartilage Thermo-Mechanobiology

We developed an *in vitro* platform allowing to recapitulate loading-induced self-heating in articular cartilage and study corresponding cell response. Our system consisted of two essential components: i) a poro-viscoelastic scaffold, with comparable dissipative capacity and equilibrium stiffness to cartilage, seeded with human chondro-progenitor cells; ii) a modular bioreactor to independently control applied mechanical loading, temperature increase as well as gas concentration and humidity levels during stimulation.

As a 3D support for cells, a fatigue resistant hydrogel (Figure 2-a,b) was developed by combination of flow-dependent and flow-independent dissipation sources to recapitulate cartilage viscoelastic properties. This was accomplished by a hybridly crosslinked (combination of covalent and physical bonds) pHEMA network and a poorly permeable macrostructure (Figure S4). Rearrangement of polymeric chains, dissociation of reversible bonds and fluid frictional drag mechanisms contributed to viscoelastic behavior of the developed hydrogel as described in detail elsewhere [14]. The stress-strain curve under cyclic loading was minimally varied showing that the developed hydrogel can preserve applied mechanical cues during multiple physiological stimulations. The poro-viscoelastic scaffold was also functionalized by RGD peptides (Figure S5) to enhance cell attachment and cell-matrix interactions during loading. The X-ray photonelectron spectroscopy (XPS) survey scan recorded on pure (non-modified) and RGD functionalized samples (Figure 2-c), confirmed the effectiveness of the RGD grafting process. As the pure pHEMA hydrogel does not contain any nitrogen atom in its molecular structure, no signal was observed in respective range for pure samples contrary to RGD functionalized samples with a peak for nitrogen at 402 eV.

**Figure 2.**
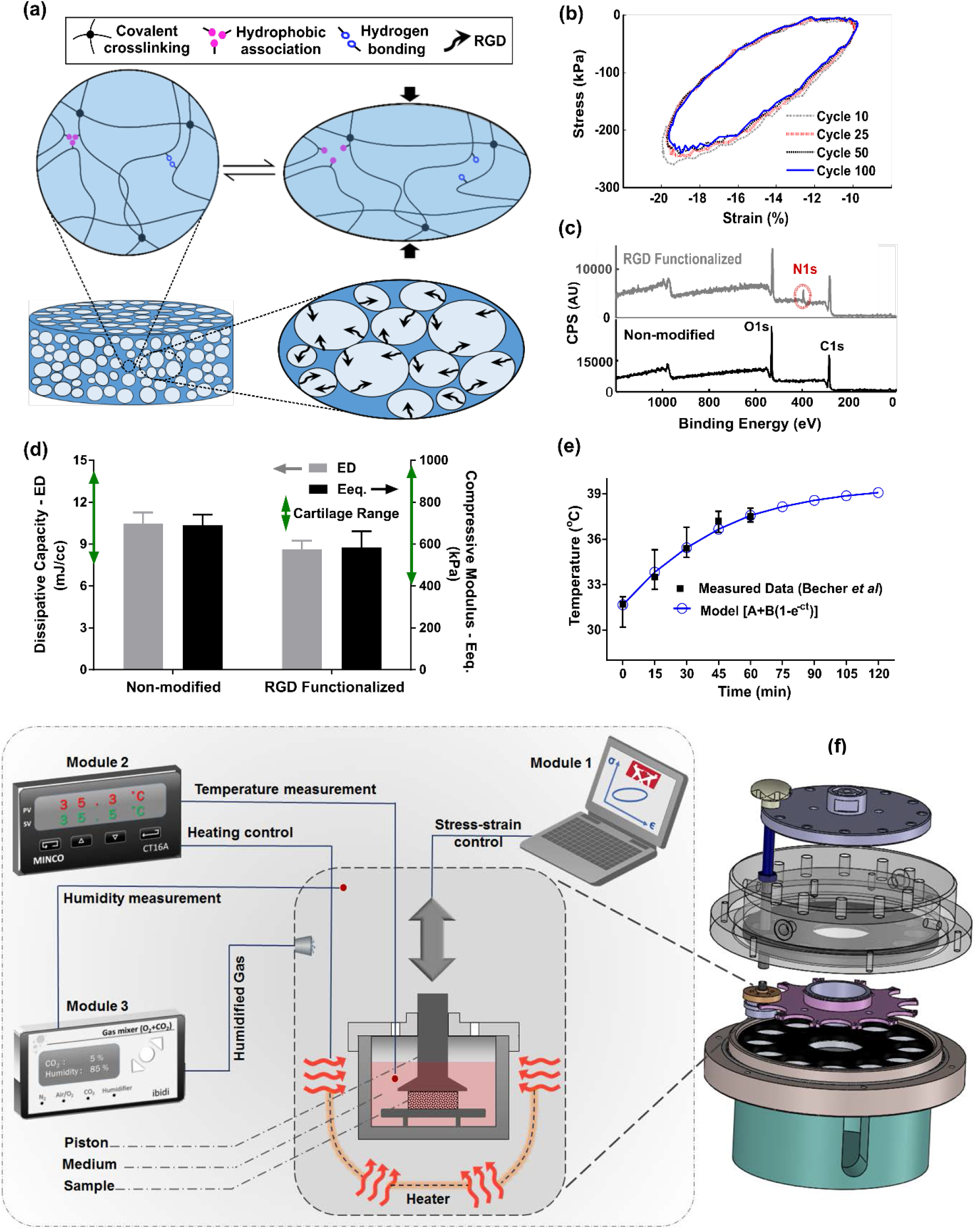
Developed *in vitro* system to study cartilage thermo-mechanobiology. (a) Poro-viscoelastic hydrogel with RGD decoration on pores. The polymeric network of the scaffold is hybridly crosslinked with physicall hydrophobic associations, hydrogen bonds and covalent crosslinks between chains. Following an applied deformation, flexible network reoraganization and fluid-solid interaction within the prorous structure occur resembling cartilage dissipative behavior. (b) The hystresis loop following loading and unloading steps was preserved over different cycles indicating fatigue resitance capability of the hydrogel. The shrinkage of the loop is mainly at the onset and remains stable after preconditioning cycles, owing to reversible sources of dissipation. (c) XPS survey scan of pHEMA porous hydrogels before and after RGD functionalization. The appearance of an N1s peak at 400 eV in the XPS spectra of functionalized samples is the evidence for a successful binding of RGD peptides to the exposed hydroxyl groups of porous pHEMA hydrogel. (d) Mechanical properties of the pure (non-modified) and RGD functionalized hydrogels and reported range of cartilage properties (green arrows) in literature. (e) The measured intra-articular temperature following jogging activity as reported in [16] and exponential fitted curve to predict the temperature evolution. (f) Modular and custom-designed bioreactor. The apparatus consists of adjustable loading system with embedded screws, chamber cap with different inlets/outlets, culture wells and pistons, wells carrier, chamber base and support. The left schematic illustrates conceptual design of the modular bioreactor for thermo-mechanical stimulation of cell-laden hydrogels.

In addition, cell proliferation in and the DNA content (Figure S6) of the functionalized constructs were significantly higher than the pure samples without RGD motifs. We could minimize the destructive effect of the bio-conjugation process on the crosslinked network and could maintain the dissipative capacity and equilibrium stiffness of the functionalized hydrogels in the range of cartilage tissue [11-13, 15] after modification (Figure 2-d). Lastly, we observed more detached cells in non-modified hydrogels without endogenous binding sites for cells when cyclic compression was combined with temperature rise (Figure S7).

Apart from tailored biomaterial properties, reliable control over applied stimuli are required to understand the effects of loading induced self-heating on chondrogenesis. Specifically, we need to simulate and control the cells thermal and/or mechanical environment during joint loading. We custom designed our bioreactor system as shown in Figure 2-f, employed an adjustable loading mechanism and applied a convection dominant heat transfer system inside wells to ensure consistent stimulation of samples. The performance of the developed bioreactor in different modes of action relevant to our study was then validated (see supplemental info for detail). Briefly, we confirmed that samples in different wells receive similar external stimuli according to the defined operation protocol. The controlled culture variables were verified by external calibrators. The temperature increase was modeled by equation 1 and its parameters were estimated based on reported *in vivo* data during jogging [16] using the least square method.

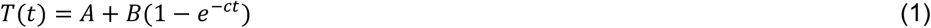

The optimally fitted temperature evolution function (A= 31.62°C, B=7.99°C and c=0.023s^-1^) predicted around 7oC temperature rise after two hours (Figure 2-e) with a good correlation coefficient (R=0.95). We controlled the evolution of the culture temperature inside the wells based on the model prediction by an external heat source to simulate intra-articular knee temperature during joint loading. Given the volume of the culture medium in each well, the size of the sample and the system boundary conditions, the accumulated lost energy from the dissipative hydrogel negligibly changed the culture temperature (<0.5°C). Our heat transfer simulation confirmed this observation when the well interface temperature was kept constant at 32°C (Figure S2). However, in an unrealistic adiabatic condition, our numerical simulation showed a temperature rise (Figure S3) due to hydrogel’s dissipative capacity (see supplemental info for detail). The isolated temperature and loading control strategy in our bioreactor design allowed us to decompose direct and indirect effects of mechanical loading on cell responses during self-heating of cartilage.

### 2.2 Temperature Evolution Following Mechanical Loading Optimally Promotes Chondrogenesis by Simulating the Cartilage Native Environment during Activity

We first evaluated viability of human chondro-progenitor cells (origin and characterization described elsewhere [35]) within the functionalized hydrogels at two different culture temperatures. The cell-seeded hydrogels were cultured for 12 days at knee intra-articular temperature at rest (32.5°C) and the conventional *in vitro* condition (37°C) corresponding to the core body temperature. We confirmed stable attachment, high viability and normal proliferation of distributed cells within the RGD functionalized hydrogels (Figure 3-a:c and Figure S6). Our results indicated that cells metabolic activity, the amount of extracted DNA and RNA (Figure S9) were slightly higher at 32.5°C compared to 37°C. These data suggest that adjusting the culture temperature at 32-33°C suits human chondro-progenitor cells metabolism which is in agreement with the reported results for human and porcine chondrocytes in monolayer and pellet cultures [19, 36].

**Figure 3.**
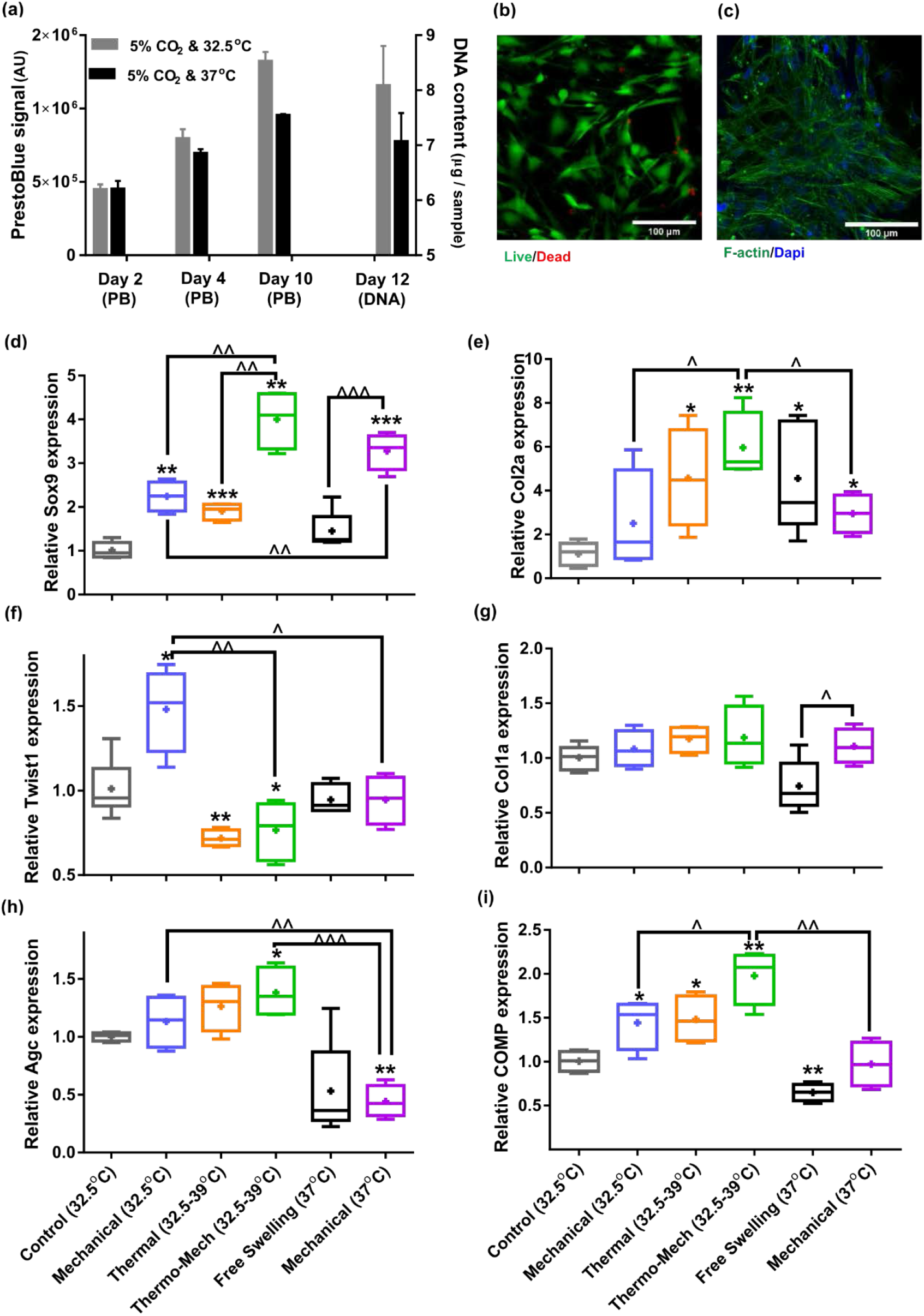
Human chondro-progenitor cells behavior inside functionalized and dissipative porous hydrogels. (a) Metabolic activity (obtained by PrestoBlue (PB) assay) and DNA content of free swelling cell-laden hydrogels in different culture temperatures corresponding to knee intra-articular temperature at rest and core body temperature (b) Live/dead labeling of cells inside porous hydrogels 4 days post seeding cultured at 32.5°C (green stain demonstrates live cells and red stain shows dead cells). (c) Actin filament and nucelous immunostaining of cells 6 days after seeding. (d:i) Comparison of the relative gene expression of cells in response to different biophysical cues applied in 3 alternate days for 90 minutes normalized to control free-swelling samples cultered at 32.5°C (significant difference with control group (^*^) or between specific groups (^^^) P<0.05; n=4-6) . A synergetic effect can be observed for Sox9 expression in thermo-mechanical stimulation group. Overal results indicate that loading-induced self heating, designated as Thermo_Mech (32.5-39°C), optimally promoted chondrogenesis.

To understand the influence of temperature and mechanical load on biological functions, gene expression data was compared by applying coupled or isolated signals. Our findings demonstrated that the loading induced self-heating (simultaneous temperature evolution and mechanical loading) positively enhanced chondrogenic responses of cells and out-performs mechanical or thermal stimulus alone (Figure 3-d:i). Importantly, we obtained an additive effect on expression of Sox9, as shown in Figure 3-d, following biomimetic thermo-mechanical stimulation designated as Thermo_Mech (32.5-39°C). Sox9 is a transcription factor and its role as one of the key regulators of chondrogenic differentiation processes is well established in the literature [37]. Upregulation of Sox9 gene under thermo-mechanical stimulation was significantly higher than the control, mechanical and thermal groups. In parallel, Twist1, which is known as an inhibitor of chondrogenesis [38], exhibited a significantly downregulated expression following the thermo-mechanical stimulation (Figure 3-f). Highest expression was also observed for functional chondrogenic markers, Col2a1, Agc and Comp, when cells received the combined physical cues. Col1a1 expression (Figure 3-g) was not significantly different between stimulated and control groups indicating enhanced chondrogenic differentiation index (Col2a1/Col1a1). Collectively, the applied thermo-mechanical stimulus provided a chondro-inductive signal by simulating the cartilage native environment under loading.

While still a promoter of chondrogenesis, isolated mechanical or thermal stimuli were not as effective as thermo-mechanical stimulus combined. Notably, application of intermittent temperature increases (from 32.5°C to 39°C during 1.5 h) on cells significantly upregulated expression of Sox9, Col2a, COMP and downregulated Twist1. Unlike traditional mechanobiological *in vitro* models with unrealistic culture temperature of 37°C, we evaluated the effect of steady-state incubation temperature on chondrogenic cellular responses. Compared to the control group, a significant fold increase was detected on expression of COMP by loading at 32.5°C. An enhanced expression of Col2a was also observed following mechanical stimulation without temperature variation. A statistically significant difference in fold increase of Sox9 was obtained between stimulated samples at 37°C and 32.5°C. This implies that chondrogenic cells respond better to compressive loading at higher temperatures which supports the common mode of action in most bioreactors for cartilage tissue engineering. Yet, simulation of loading induced self-heating condition better promoted chondrogenic markers compared to mechanical stimulus at constant temperatures (32.5 or 37°C). This indicates that cells could sense the transient temperature rise. The transient effect should therefore be considered as a potential potent regulatory cue besides external stress.

### 2.3 Cartilage Thermo-mechanotransduction Is a TRPV4-mediated and Calcium Dependent Signaling Mechanism

Knowing that oscillation of intracellular calcium is an important regulator of chondrogenesis, the role of Ca^2+^ signaling in thermo-mechanotransduction process was evaluated. We also examined the contribution of multimodal TRPV4 ion channels to this process as a potent Ca^2+^modulator (Figure 4-a). In this regard, we initially assessed the presence and functionality of TRPV4 channels on human chondro-progenitor cells. Immunostaining of cells using a specific antibody confirmed the presence of TRPV4 channels on cells (Figure 4-b) seeded in functionalized hydrogels. The functioning of TRPV4 was confirmed by live florescence imaging of cells following administration of 10 nM TRPV4 agonist, GSK1016790 (GSK101). We could detect multiple dynamic peaks after GSK101 delivery (following 70 seconds baseline recording) when analyzing intracellular calcium signals as shown in Figure 4-c for 3 representative cells (see supplemental movie 1 for whole response of a group of cells). We then verified significant inhibition of TRPV4-mediated calcium signaling (see supplemental movie 2) by using 10 µM TRPV4 antagonist (GSK205) in culture medium one hour before imaging. Moreover, full chelation of extracellular calcium sources was obtained by employing 2 mM EGTA in the culture medium which completely diminished cytosolic calcium variation. We also evaluated the cell viability in presence of effective quantities of employed chemicals (DMSO as carrier, GSK101, GSK205 and EGTA as agonist/antagonist) in our pathway study. Results of the assay (Figure 4-d) for all tested conditions, showed more than 85% viability at day 8 after using chemicals in culture medium for 4 hours during two different days.

**Figure 4.**
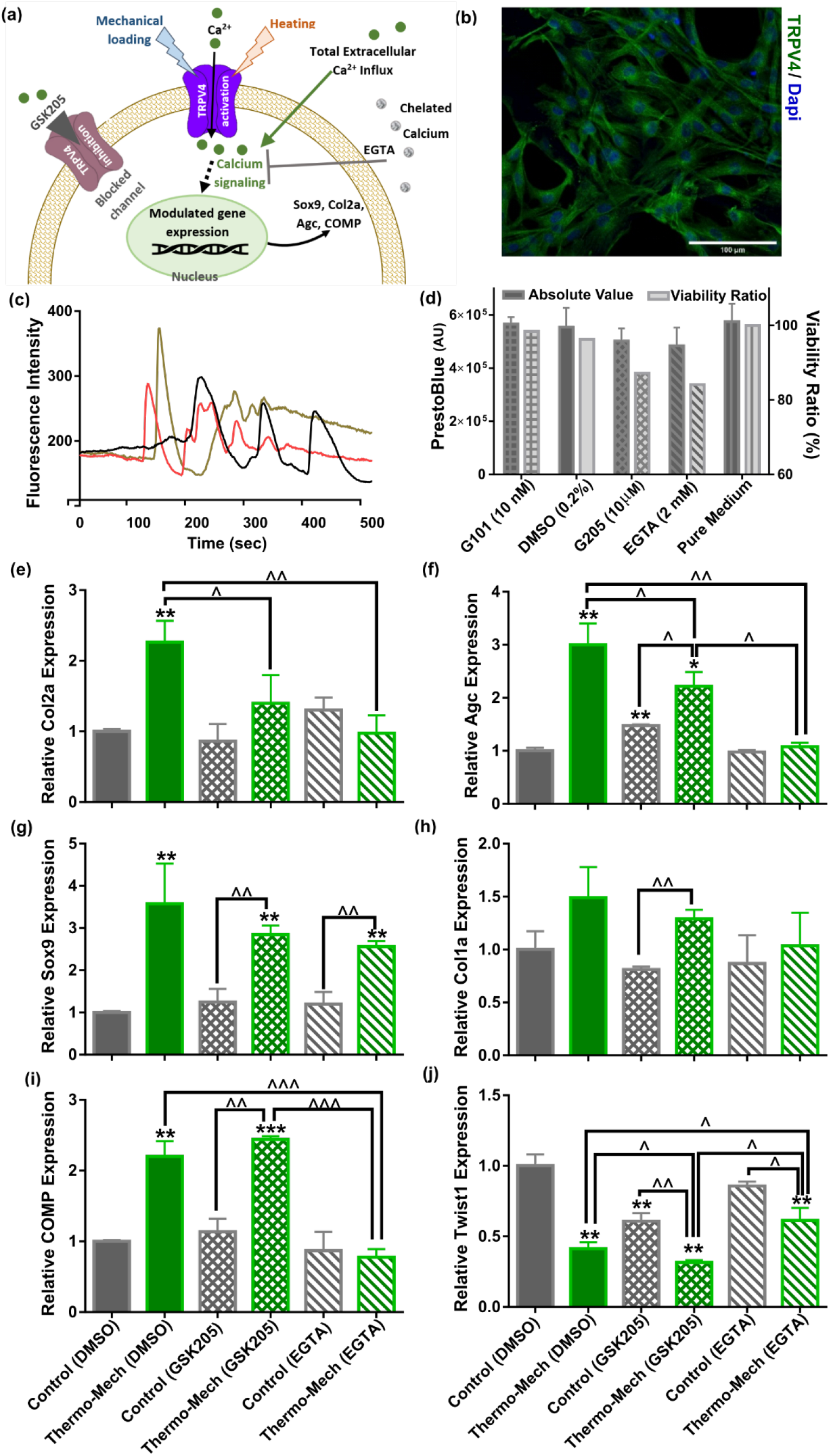
Calcium signaling mechanism, functional charactrization of TRPV4 ion channels expression in human chondro-progenitor cells and gene expression results with maniplualted pathways regulating cytosolic Ca^2+^variation. (a) Schematic of TRPV4-mediated calcium signaling for transduction of thermo-mechanical cues which could be inhibited by GSK205 (TRPV4 antagonist). Generally, the interacellular calcium can be varied through different pathways and all of them are affected when free Ca^2+^ ions are chelated with EGTA. (b) Immunostaining of TRPV4 channels. Successful binding of the specific-TRPV4 antibody to human chondro-progenitor cells confirmed expression of TRPV4 channels on cell-laden hydrogels. (c) Analysis of cytosolic calcium intensity over time by flouresence imaging. Response of cells to TRPV4 agonist confirmed functionality of the gates for modulation of interacellular calcium content. (d) Viability of cells in presence of pertinent agonsit/antagonist of calcium pathways. (e:j) Relative gene expression of cells in different groups normalized to free-swelling control group with DMSO as drug carrier when employing specific antagonist for inhibiting TRPV4 channels (GSK205) or chelating extracellular calcium sources (EGTA). The control groups are free-swelling samples cultured at 32.5°C and stimulated samples receiveing combined thermo-mechanical cues at the same culture medium containing DMSO, GSK205 or EGTA (^*^ significant difference with control group with P<0.05; ^^^ significant differences between specific groups with P<0.05; n=3-4). Extracted results indicated that thermo-mechanotransduction signalling cascade was almost incomplete without extracellular calcium sources and TRPV4 channels played a key role in translation of perceived physical cues to calcium transients.

The inhibition of TRPV4 channels with 10 µm GSK205 diminished the elevated expression of the main chondrogenic markers when cells received the combined physical cues. As shown in Figure 4-e:h, a significant downregulation for expression of Col2a and Agc genes was obtained in stimulated samples with and without TRPV4 antagonist. We also observed a slight decrease in Sox9 and Col1a expression with blocked TRPV4 channels. Inhibiting TRPV4 activity in free-swelling samples did not affect relative expression of the Col2a, Sox9, Col1a, and Comp (Figure 4-i). In contrast, we could detect a significant downregulation of Twist1 in control samples with GSK205 (Figure 4-j). Consequently, further downregulation was obtained for Twist1 gene by thermo-mechanical stimulation when TRPV4 activity was inhibited. Our results also indicated that TRPV4 acts as signal integrator for thermal and mechanical stimuli (Figure S10). By comparing the Col2a and Sox9 gene expression for either cue alone, with and without GSK205, one can observe reduced levels of these markers when TRPV4 is inhibited. The downregulation in blocked TRPV4 channels condition is statistically significant for Sox9 only in case of mechanical loading and for Col2a only in case of thermal stimulus.

Without application of external physical stimulus, we could not observe a significant change on relative gene expression of cells in presence of 2 mM EGTA. Similarly, the promoted expression of chondrogenic markers following thermo-mechanical stimulation was almost abrogated without external Ca^2+^sources. In particular, we measured a significant reduction in expression of Col2a, Comp and Agc genes and a significant increase in expression of Twist 1 (Figure 4-e,f,i & j). The average Sox9 expression (Figure 4-g) was also decreased; however, the downregulation was not statistically significant in the absence of external Ca^2+^. Taken together, our results suggest that TRPV4 mediates thermo-mechanotransduction processes and the Ca2^+^ signaling participates more in modulation of functional ECM markers (collagen type 2, cartilage oligomeric matrix protein and aggrecan genes) than Sox9 transcription factor.

## 3 Discussion

Many groups, including ours, have already demonstrated the positive impact of mechanical loading on chondrogenesis in explant and cell-scaffold constructs [5, 7, 8, 14, 39-41]. However, previous *in vitro* mechanobiological studies typically considered a constant culture temperature of 37°C which does not correspond to the physiological *in vivo* condition in human knee. For the first time, we investigated cartilage thermo-mechanobiology and demonstrated the synergetic effect of heat and loading stimuli at the transcriptional level of chondrogenesis. We further demonstrated that the thermo-mechanotransduction is indeed calcium dependent and TRPV gates on the plasma membrane play a key role in this process. This study also revealed that decomposed thermal cue, as a by-product of loading, could itself promote chondrogenesis similar to or even better than the mechanical signal alone. Our findings support the concept that ECM viscoelasticity can indirectly influence biological response of cells as self-heating phenomenon links the temperature evolution to cyclic loading of viscoelastic materials.

### 3.1 Simultaneous Consideration of Inter-related Cues for Cartilage Thermo-Mechanobiology

As already well established, the biophysical cues significantly influenced the cellular responses in musculoskeletal tissues [1-3, 6]. In a mechanistic view point, structural and material properties of ECM microenvironment are acting in association with external stresses to generate physical cues for cells [3, 9, 10]. The cartilage dissipative capacity originates from intrinsic matrix viscoelasticity and solid-fluid interaction under an applied loading. These dissipative sources are associated with the matrix deformation, hydrostatic pressure and interstitial fluid flow around cells which were previously considered as mechanobiological regulators [4, 7, 8]. Cartilage dissipation, therefore, could be a functional property for chondrogenesis overarching all solid and fluid related mechanical signals [13, 14]. In parallel, energy dissipation in avascular cartilage during joint loading generates heat and causes the temperature evolution over time [15]. The absolute temperature rise in cartilage microenvironment also depends on the joint boundary condition (e.g. the ambient temperature and heat flux by surroundings tissues). Regardless of the heat source(s) and of the complex heat transfer mechanisms inside articular cavity, the varied temperature during joint loading could affect chondrocyte biological responses. To simulate such conditions and study cartilage thermo-mechanobiology, we developed a fatigue resistant dissipative hydrogel for 3D culture of cells and a modular bioreactor to apply respective biophysical cues *in vitro* as there was not an available platform for this purpose elsewhere.

The reciprocal relationship between cells and ECM governs cartilage hemostasis as chondrocytes receive physical cues and regulate tissue turnover to maintain its functional characteristics (e.g. dissipation). An altered gene expression is generally considered as the first indicator for cells responsiveness to conveyed cues which could ultimately lead to long term matrix deposition depending on other contributing factors. Collagen type II (Col2a), aggrecan (Agc) and SRY-related HMG-box gene 9 (Sox9) genes are the most utilized chondrogenic markers at the molecular level for mechanobiological studies [7, 41]. Not only does Sox9 contribute to early chondrogenesis and matrix synthesis [37], but it also inhibits chondrocyte terminal differentiation [42]. Col2a and Agc are anabolic markers of main ECM components in cartilage tissue and COMP gene controls synthesis of the linkage protein with multiple binding possibilities to integrate and stabilize the ECM [43]. Twist-related protein 1 (Twist1) is known to hinder chondrogenesis by direct inhibition of Sox9 as the master regulator of the process [38]. Our customized model system enabled us to show that chondrogenic cells immediately responded to biomimetic thermo-mechanical cues by altering their gene expression and that cells are able to sense the transient temperature rise. We evaluated cellular reactions after three rounds of stimulation to consider their probable desensitization and estimate a robust transcriptional response. However, given the transient nature of the chondrogenic markers to an applied stimulation [44], analysis of results at different time points could help to better understand the gene expression profile. Additionally, it still remains to determine to what extend the promoted chondrogenic gene expression could enhance downstream protein expression and functional properties.

### 3.2 Cartilage Thermo-mechanics Portrays New Opportunity to Advance Chondrogenesis *in Vitro*

The expression of chondrogenic markers was enhanced by the applied physical stimulus in all groups except for Agc and COMP in one group (Figure 3).The obtained gene expression data for Col2a1, Agc, Col1a, Sox9, Twist1 and COMP clearly show that thermo-mechanical stimulus generates more potent signaling to elicit cell responses compared to either cue alone. One compelling aspect of this finding is that employing bioreactors which are capable of controlling temperature evolution during dynamic loading, should be preferred for tissue engineering applications. Indeed, chondro-inductivity can be enhanced by the condition which simulates the *in vivo* cartilage more similarly as shown by other multi-functional bioreactors [7, 39]. For instance, simulating complex physiological loading conditions in articulating joints by combined compression, sliding and shear was shown to be more effective than a single loading mode [39]. In addition, combining hypoxia culture with multi-directional loading performed better than normaxia [45] as low oxygen level could direct chondrogenesis. Accordingly, investigating controlled oxygen tension combined with thermo-mechanical stimulation might shed more light on the *in vivo* chondrogenic process.

Dynamic temperature evolution also had promising impact on transcription of chondrogenic markers in the current study where applied thermal stimulus out performs pure mechanical loading. Significant upregulation in Col2a1, COMP and Sox9 expression as well as downregulation of Twist1 gene clearly indicate enhanced cellular responses to isolated self-heating cue (temperature rise without loading). In line with our findings, available evidence in the literature suggests that proper heat stimulation can enhance and accelerate chondrogenic differentiation processes [18]. Over stimulation by high temperature, however, reduces the cell viability and inhibits cartilage matrix synthesis [19]. Therefore, it is important to simulate the dynamics of thermal environment *in vivo* and optimize the influence of thermal stimuli by defining a biomimetic intermittent temperature rise. Considering clinical translation, it is encouraging to have chondro-inductive cues only with application of careful heating to simulate joint thermal environment during physical activity. Such stimulus can be simply delivered to patients in need by development of a customized heat therapy device for joint disease. It remains to elucidate if this argument is true for osteoarthritic cartilage as damaged chondrocytes may respond differently to biophysical cues as reported recently [46].

### 3.3 Calcium Signaling Is a Major Mechanism of Action in Transduction of Thermo-mechanical Cues Sensed by TRPV4 Channels

Understanding the translation mechanisms of external stimuli and their interaction with molecular signaling pathways is essential for development of therapeutic products to modulate the target regulators. It is well established that TRPV4 is a key mediator in transduction of mechanical signals in chondrocytes which is necessary for maintenance of cartilage tissue [5, 26, 46]. In parallel, the reported activation temperature around 30oC for TRPV4 [26, 33] makes this channel a potential mediator of loading-induced self-heating in cartilage. Our study revealed that TRPV4 is a potent modulator for expression of Col2a and Agc genes in response to thermo–mechanical cues. These results are in agreement with the data reported for post transcriptional markers determined by protein expression and functional properties of cell-encapsulated hydrogels under mechanical stimulation [5, 40]. The obtained gene expression profile of Sox9 in this study suggested a partial contribution of TRPV4 to its transcription in response to thermo-mechanical cues. This means that there are parallel mechanisms for sensing the applied stimuli and eliciting Sox9 enhancement which could be other transmembrane channels, primary cilium or integrin [7, 9, 24, 29]. Integrin-mediated pathways arising from cell-matrix interaction are mechanisms by which cells may sense the mechanical cues in the surrounding microenvironment [3, 20]. Since RGD motifs were conjugated onto the pores of the developed hydrogels, a stable and effective cell-matrix interaction during loading can be expected. According to our results, significant upregulation of Sox9 in the mechanically loaded samples with and without GSK205 (Figure S10) could imply the contribution of an integrin pathway. Of note, previous studies also showed that integrin could regulate activation of ion channels in chondrocytes and cell-matrix interactions could modulate calcium signaling dynamics in response to physical cues [22, 25, 47]. We could not find any direct thermo and/or mechano-biological study in the literature reporting even partial effect of TRPV4 blocking on Sox9 expression in chondrogenic cells. However, in agreement with our findings, *in vivo* and *in vitro* studies have shown that pharmacological activation/inhibition of TRPV4 channels could mediate Sox9 expression [31, 48].

TRPV4 activity can modulate intracellular calcium transients and literature findings indicate that a well-adjusted TRPV4-mediated Ca^2+^ signaling is necessary for correct biological functioning of chondrocytes [5, 29-32]. We found that full removal of external calcium sources could further diminish the enhanced chondrogenic response obtained by thermo-mechanical cues when compared with TRPV4 inhibition alone. This behavior was consistent in gene expression profiles of the investigated markers and induced transcription returned toward control levels in the presence of EGTA. These findings indicate that calcium is a major signaling mechanism for thermo-mechanotransduction. Intracellular calcium oscillation is a key signaling mechanism in chondrocytes and can be triggered by different types of mechanical stimuli such as strain [24, 25] and fluid flow [22]. Additionally, the dynamic characteristics of calcium signaling (e.g. amplitude and duration of Ca^2+^ transients) were shown to be temperature dependent [25]. Our results suggest that the signaling cascade of thermo-mechanotransduction process is almost incomplete without external calcium sources. Previous studies also showed that extracellular Ca^2+^ source is necessary for calcium signaling processes and increase in cytosolic Ca^2+^ is not possible without extracellular Ca^2+^, even when intracellular stores are full of calcium [23, 24].

When thermo-mechanical cues are applied, downregulation of Agc and COMP and upregulation of Twist1 were significantly different in the presence of EGTA compared to when TRPV4 antagonist was used (see Figure 4). This different response could arise from two plausible reasons. First, TRPV4 channels were not fully blocked by GSK 205 and the conveyed signal was strong enough to partially activate some of the channels. This is consistent with other studies [21, 24] where quantitative results indicated that 10 µm GSK205 significantly inhibited but did not fully abolish TRPV4 channel activity. A second possibility would be that, parallel thermo-mechanically responsive pathways besides TRPV4 were involved in Ca^2+^ mediation in order to make this process more effective and robust. This is not surprising as accumulating evidence in the literature suggests that distinct calcium pathways, including stretch-activated channels (e.g. PIEZO1), voltage-gated calcium channels (e.g., T-type VGCC), purinergic receptors (e.g., P2Y or P2X), PLC-IP3 induced endoplasmic reticulum and TRP family channels (e.g., TRPV3), are directly or indirectly influenced by an externally applied stimulus [23, 24, 28, 29]. Determining other probable contributing mediators in thermo-mechanotransduction processes are thus important next steps and need further investigation.

In summary, different aspects involved in cartilage thermo-mechanobiology following joint loading were analyzed by developing a novel customized *in vitro* model. Our findings have demonstrated the superior effect of thermo-mechanical cues on chondrogenesis compared to either cue alone. Indeed, the applied synergistic stimulus provided a chondro-inductive signal by simulating the cartilage native environment under loading. This study also has shown that biomimetic temperature evolution as by-product of mechanical loading could itself induce biological responses of cells. Moreover, TRPV4 ion channels were identified as a key mediator of thermo-mechanotransduction process. Thermo-mechano-responsive nature of TRPV4 channels makes them a key target for future investigation on thermo-mechanobiology. We found calcium signaling as a contributing mechanism of action translating the effect of the externally applied cues to intracellular biochemical signals for cells. However, other pathways are also playing parallel roles, especially for Sox9 transcription, which therefore would necessitate further understanding of thermo-mechano-transduction processes overall.

## 4 Method

### 4.1 In vitro culture and biological evaluation

Cell growth medium consisted of DMEM containing L-Glutamine, 4,5 g/l D-Glucose and Sodium pyruvate, (Life Technologies) which was supplemented with 10% FBS (Sigma) and 1% L-Glutamine and Penicillin (Life Technologies). Cells were grown to 90% confluence, trypsinized and redistributed again in 2D culture up to passage 5 before seeding into scaffolds (∼1.2 mio/scaffold). Stimulation medium was prepared with cell growth medium without FBS but having 10% ITS IV (Life Technologies) and 1% Vitamin C. MTT staining Kit I (Roche) was used to control the cell distribution and Viability Assay Kit (Biotium) was utilized for live/dead evaluation according to the manufacturer protocols. By using the MTT reagent, the purple formazan is produced due to the cells metabolic activity and therefore reveals cell distribution within the hydrogels. Viability assay was performed via a cell-permeable dye (calcine) for staining live cells and a cell-impermeable dye (ethidium homodimer) for staining dead cells having a damaged cell membrane. The Hoechst 33258 dye (ThermoFisher Scientific) was employed for DNA quantification to evaluate the impact of hydrogel functionalization on cell attachment. Briefly, the pure and RGD-modified hydrogels (with and without cells) were cut in small pieces and incubated overnight inside 500 ml papain digestion buffer at 65°C. Then, by dilution of 10 µl of digested solution in 140 µl of the dye (0.2 µg/ml), the emission signal of samples was measured at 460 nm by a Wallac microplate reader after excitation at 355 nm. The DNA content of the samples was finally determined by using a standard curve extracted from sequential dilutions of Calf Thymus DNA (ThermoFisher Scientific) as calibrators. Prestoblue viability kit (ThermoFisher Scientific) was also used to monitor the cell proliferation inside the hydrogels during culture period [49] based on manufacturer protocol. Briefly, the Prestoblue reagent was 10 times diluted in culture medium and samples were incubated inside. The fluorescent signal was then measured with Wallac microplate reader for excitation/emission wavelengths of 544/590 nm, respectively.

### 4.2 Design of Modular bioreactor for thermo-mechanobiological study

The bioreactor chamber was custom designed and supplemented with different modules to reliably alter dynamic culture conditions. The mechanical design of the bioreactor chamber was optimized to ensure sterility and application of reproducible spatiotemporal stimuli on samples. The chamber, as shown in Figure 2-a, contains 12 circularly arranged cylindrical wells for culture of cell-laden hydrogels. The set-up is compatible with the Instron uniaxial loading machine (Instron-E3000, Massachusetts, USA) to apply the mechanical loading (Figure S1) according to reported values for knee cartilage deformation during walking [50]. The bioreactor was fixed to the Instron apparatus at its base by a support, and was stimulated from above by a set of pistons attached to the actuator. To control culture temperature, a thermo-foil heater (Minco-HM6975) was embedded below an aluminum conductive disc supporting culture wells. The temperature was measured by a miniature RTD sensor (Minco-S308) placed inside one of the wells. The evolution of culture temperature was feedback controlled by a PID microprocessor (Minco-CT16A, Minnesota, USA) based on the error signal between the desired and monitored temperatures. The parameters of the PID controller were tuned experimentally based on the standard Ziegler-Nichols method. The heat controller was tuned to regulate the culture temperature defined by sequential ramps with different slopes according to the model prediction. This strategy was employed to simulate the temperature increase (occurring in knee joint with its irregular boundary conditions due to cartilage self-heating) in our small-size scaffolds (Ø:8, t: 2.2 mm) surrounded by the relatively large volume of culture medium (∼2 mL) inside the bioreactor wells. To minimize the temperature gradient due to conduction within the sample from bottom to top, a medical graded thermoplastic part (PPSU) was incorporated within the stainless-steel wells. The geometry of PPSU part was carefully designed to make convection the dominant heat transfer mechanism to samples. A flexible silicon support was also employed to keep the position of samples on the center during dynamic loading. To maintain sterility, the wells were isolated from the outside environment with perforated solid caps covered with gas permeable filters (OpSite Flexigrid, Smith &Nephew, Hull, UK). The CO_2_/O_2_ concentration and humidity inside the chamber were regulated by an external gas mixer (ibidi-Gas Incubation System, Martinsried, Germany) injecting humidified gas into the bioreactor. The humidity and gas concentration was directly measured inside the chamber with external calibrator probes to ensure correct functioning of the gas regulator, humidifier and embedded sensors of the device.

### 4.3 Dissipative hydrogel fabrication and mechanical characterization

Salt leaching method was employed for porous hydrogel fabrication by polymerization of HEMA-EGDMA precursor (4.8% molar ratio) inside molds containing sieved salt particles (150-250 µm) as described previously [14, 51]. Energy dissipation was determined by calculating the hysteresis area of stress-strain curve during loading and unloading of cycle 100. Equilibrium Young modulus (Eeq.) was measured by application of sequential stress relaxations and finding the slope of the corresponding relaxed stress in range of 10-20% strain values.

### 4.4 RGD functionalization of hydrogels

The functionalization of pHEMA porous hydrogels with RGD peptide were performed following a published protocol with slight change [52] (Figure S5). The pendant hydroxyl groups of pHEMA hydrogels were firstly activated by 4-nitrophenyl chloroformate (NPC) and then RGD peptides were conjugated. For this purpose, lyophilized hydrogels were vacuum-swelled in the freshly prepared activation mixture of 1.41 g NPC in 15 ml acetone and 1 ml distilled triethylamine. The activation process was continued by agitation of samples in the solution for 1 hour at room temperature. The samples were then extensively rinsed with acetone and methanol several times to remove excess NPC. Afterwards, the NPC-activated samples were transferred to the methanol solution containing 1 mM RGD peptides and 2 mM 4-dimethylamino-pyridine. To complete the functionalization process, the samples were incubated overnight at room temperature under gentle shaking in the reactor covered with aluminum foil. The RGD functionalized samples were then rinsed three times with pure methanol for 2 minutes. To ensure removal of any non-reacted NPC group, the samples were incubated in 0.5 M solution of ethanolamine in methanol for 20 minutes under gentle shaking. The samples were finally rinsed several times with methanol and bi-distillated water to wash non-specifically adsorbed peptides and entrapped NPC molecules inside the porous hydrogels.

### 4.5 Simulation of heat transfer inside bioreactor culture well

To ensure independent control over applied loading and temperature evolution in bioreactor culture wells, we developed a finite element (FE) heat transfer model using COMSOL Multiphysics software (Burlington, MA, USA). The hydrogel dissipation was considered as a heat source generating 9000 W/m^3^(determined experimentally based on area of hysteresis loop at 1Hz) in each step of cyclic compression. The solid and the fluid phases of the porous scaffold were coupled based on theory of poro-elasticity and a linear visco-elastic behavior for the solid part was assumed as described in our previous work [14]. Moreover, the heat transfer module was coupled to the resultant pressure/velocity field in porous media to consider convective heat transfer mechanism. In the model, the 2D axisymmetric cross section of cylindrical samples (2.2 mm thickness and 4 mm radius) and culture medium were divided to finite elements and boundary conditions were applied as shown in Figure S2 and S3. The cyclic displacement (10% strain @ 1 Hz) was applied on the interface of the piston with sample-medium domains and stress, strain, velocity, pressure and temperature fields were solved in a fully coupled manner. The model was examined for different mesh sizes to ensure that results are not changed with elements size. We assumed isotropic mechanical behavior for the poro-viscoleastic scaffold with material properties were assigned according to Table S2.

### 4.6 Cartilage loading induced self-heating *in vitro*

Human epiphyseal chondro-progenitor cells were expanded in 2D culture according to standard cell culture protocols [35] before seeding them into porous hydrogels. Hydrogels were disinfected by graded ethanol. Cells (∼1.25 million per sample) were infused into them by an optimized compression released induced suction method [53]. The effect of the different biophysical stimuli was then assessed by application of respective signals on cell-laden hydrogels. The reported *in vivo* data in the literature for deformation of tibio-femoral cartilage during walking [50] and temperature rise during jogging [16] activities were used to apply respective stimulus *in vitro*. After the seeding step, all samples were pre-cultured for 4 days in cell growth medium inside standard incubators (32°C or 37°C and 5% CO_2_). After this step, a cell stimulation medium was used and the thermal (32 to 39°C), mechanical (10% pre-strain, 10% amplitude at 1 Hz) or thermo-mechanical (their combination) stimulation were applied on samples starting from day 6. Three intermittent stimulations in alternate days were applied on the treated samples for 90 minutes while the control group was cultured in equivalent conditions but without receiving any stimulus (Figure S8).

### 4.7 RNA isolation and real-time quantitative RT–PCR

Total RNA was extracted using the NucleoSpin® RNA (Macherey-Nagel) after preparation steps. Briefly, samples were put in a 2 ml Eppendorf tube containing 300 µl Trizol. Hydrogels were disrupted with a polytron (Kinematica AG, Switzerland), while keeping them cold on dry ice. Then, 100 µl chloroform was added and samples were centrifuged for 5 minutes at 12000 rpm at 4°C. The aqueous phase was transferred to 2 ml phase lock tubes and centrifuged for an additional 5 minutes at 12000 rpm at 4°C. The aqueous phase was carefully transferred to 1.5 ml Eppendorf tubes and the extraction was completed by adding a RNA carrier and following the XS kit protocol. The RNA was quantified using the Nanodrop Lite Spectrophotometer (Thermo Scientific) and reverse transcription of 1 µg RNA was carried out using Taqman® Reverse Transcription Reagents (Applied Biosystems, Massachusetts, USA). Fast SYBR® Green Master Mix (Applied Biosystems) was used for PCR amplification in a final volume of 20 μl containing 10 ng of synthesized cDNA. The PCR amplification was performed for each sample by StepOnePlus Real-Time PCR device (Applied Biosystems) using specific primers for different genes (Table 1). The cycling steps were defined as an initial 95°C for 2 min followed by 40 cycles of amplification. Gene expression data were analyzed using the comparative ΔΔCt [54] method with RLP13a as the reference gene [55].

### 4.8 Immunostaining

Characterization of TRPV4 protein on human chondro-progenitor cells was conducted by employing an anti-TRPV4 antibody (ab191580, Abcam) via standard immunostaining technique. Briefly, samples were fixed by PAF (4%) and cells were permeabilized by triton (0.25%). After rinsing by PBST, samples were incubated for 1 hour in blocking solution (1% BSA) and then overnight in TRPV4 specific antibody (5ug/ml). Finally, the secondary antibody (goat-anti rabbit) conjugated with Alexa flour 488 was used to visualize TRPV4 channels by a confocal fluorescent microscope (LSM 700-Zeiss).

### 4.9 Calcium Imaging

Calcium imaging was performed by staining samples with 5 µM Flou 4-AM reagent (F14201, Thermofisher) according to manufacturer protocol. Activation of TRPV4 channels was assessed by using 10 nM GSK101 in HBSS solution. The samples were imaged by spinning disc confocal microscopy (Visitron) up to 10 min and the effect of TRPV4 agonist was analyzed. To diminish total extracellular calcium and inhibit TRPV4 mediated calcium influx, free calcium ions were chelated with 2 mM EGTA and blocked the channel with 10 µM TRPV4 antagonist, GSK205, respectively.

### 4.10 TRPV4 pathway and calcium signaling study

Gene expression of cell-laden hydrogels in different groups was tested by using medium containing vehicular control, TRPV4 antagonist and calcium chelator. The samples were stimulated according to our standard thermo-mechanobiological experiment protocol (Figure S8) while the medium was supplemented with 0.2% DMSO as control vehicle, 10 µM GSk205, or 2 mM EGTA. These chemicals were added to the culture wells 1 h before start of stimulation and removed half an hour after stimulation to limit exposure of cells to pathway antagonists. Afterwards, the samples were washed twice by fresh medium and stimulation medium was added without chemicals.

### 4.11 Statistical Analysis

All biological and mechanical experiments were analyzed with at least three replicates per test condition. The statistical significance between different study groups was determined by Student t-test (Welch assumption). Statistical comparisons were performed with GraphPad Prism software (GraphPad). Calcium imaging evaluations were performed at different time points each covering more than 50 cells per test condition during TRPV4 activation, inhibition and Ca^2+^ chelation.

## Supporting information

Supplemental information for main manuscript

## Acknowledgements

We thank Sandra Jaccoud and Dr. Nathalie Burri for their technical support. We acknowledge the contribution of the Polymers Laboratory of EPFL and in particular Prof. Harm Anton Klok for his advice on functionalization of pHEMA hydrogels. We also appreciate EPFL mechanics workshop and in particular Marc Jeanneret for his precious technical support in design of the bioreactor chamber. This work was supported by the Swiss National Science Foundation (#310030_149969/1 and #CR23I3_159301).

## Author contributions

N.N. designed the customized model system and performed the in vitro tests. N.N., and D.P.P. conceptualized the study and designed the experiments. J.W. and N.N. developed the scaffolds functionalization method. L.B. and N.N. executed mechanical characterization and validation tests. Y.G. and N.N. conducted calcium imaging and heat transfer numerical simulations. N.N., P.K., P.A.S, and D.P.P. contributed to the analysis of the results. L.B., P.K. and N.N designed and prepared the figures.

N.N. wrote the manuscript and P.A.S., L.A., D.P.P., L.B., and P.K. reviewed the manuscript. D.P.P. supervised the project.

## Supplementary info

Supplemental data accompanies this manuscript as a separate file.

