## Supplemental information for main manuscript for "Temperature Evolution Following Joint Loading Promotes Chondrogenesis by Synergistic Cues via Calcium Signaling"

EPFL/STI/IBI/LBO, Station 9, 1015 Lausanne, Switzerland

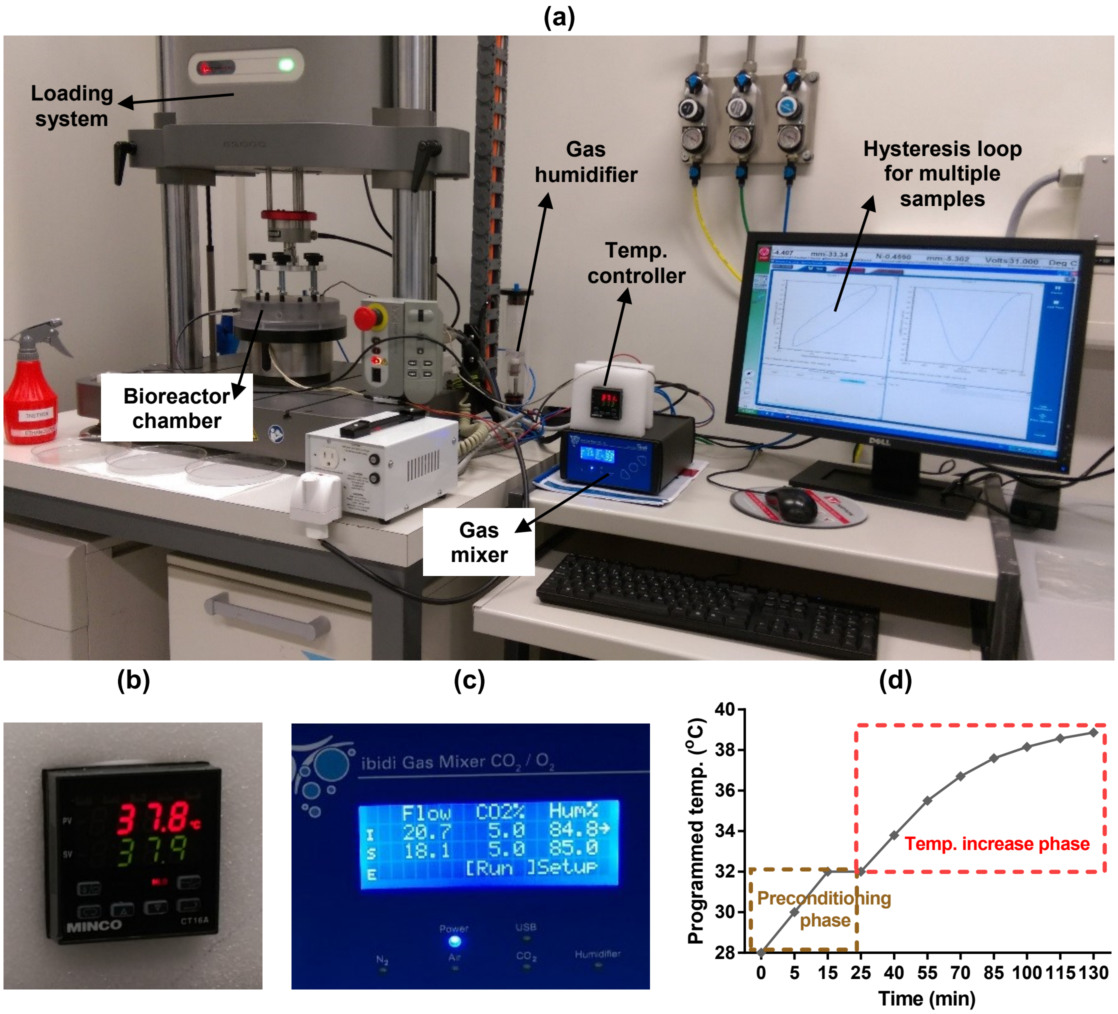


**Figure S1.** Functions evaluation of the developed bioreactor for simulation of cartilage loading-induced self-heating. (a) Validation of the performance of the developed device for thermo-mechanical stimulation. Modular design of the bioreactor permitted independent control over applied mechanical load, culture temperature, $CO_2/O_2$ gas concentration and humidity level. The Instron device was programmed to apply controlled cyclic strain on samples according the reported values in the literature for deformation of knee cartilage during stance phase of walking (7-23%) at 1 Hz frequency. Smooth curve of hysteresis loop shows that samples are uniformly loaded and deloaded. (b) The culture temperature closly track the programed temperature based on the temperature evolution model. (c) The gas mixer was set on normaxia condition to regulate standard 5% $CO_2$ and 85% humidity levels inside the chamber. (d) Simulation of the temperature increase during cyclic compression with the developed bioreactor after a preconditioning phase. During the preconditioning, the temperature of the medium inside the culture wells reached 32°C after 10-15 minutes to simulate the cartilage temperature at rest. Meanwhile, the humidified gas injection provided a stable 5% $CO_2$ and 85% humidity inside the chamber. After equilibrating the system, the mechanical stimulation began and the temperature could either be changed to evolve according to the prediction of the curve fitted data or be kept constant during the cyclic compression period. The error between desired and realized temperatures was less than 0.5°C when the PID coefficients were optimized. It was also verified that actual recorded temperature inside the wells in different positions were able to closely follow the programmed regime with a maximum error of 0.5°C.

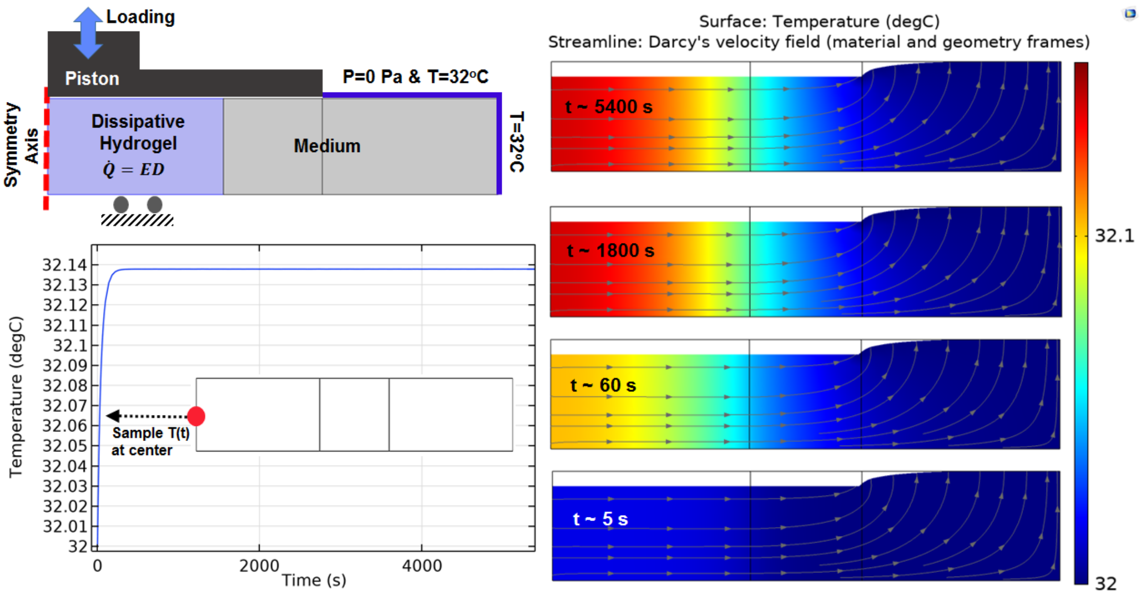


**Figure S2.** Heat transfer simulation in one culture well of the bioreactor by constant temperature buondary condition (32°C) at outer interface. The temperatue of the sample and medium was minimally varied over stimulation period from initial value ( 32°C) to maximum 32.14°C in core of the sample. The color map illustrates temperature and the arrowed curves are velocity streamlines.The obtained results confirm that the external heat supply controls the culture wells temperature independent from the applied mechanical loading. The isolated temperature and loading control strategy in our bioreactor design allowed us to decompose direct and indirect effects of mechanical loading on cell responses during self-heating of cartilage.

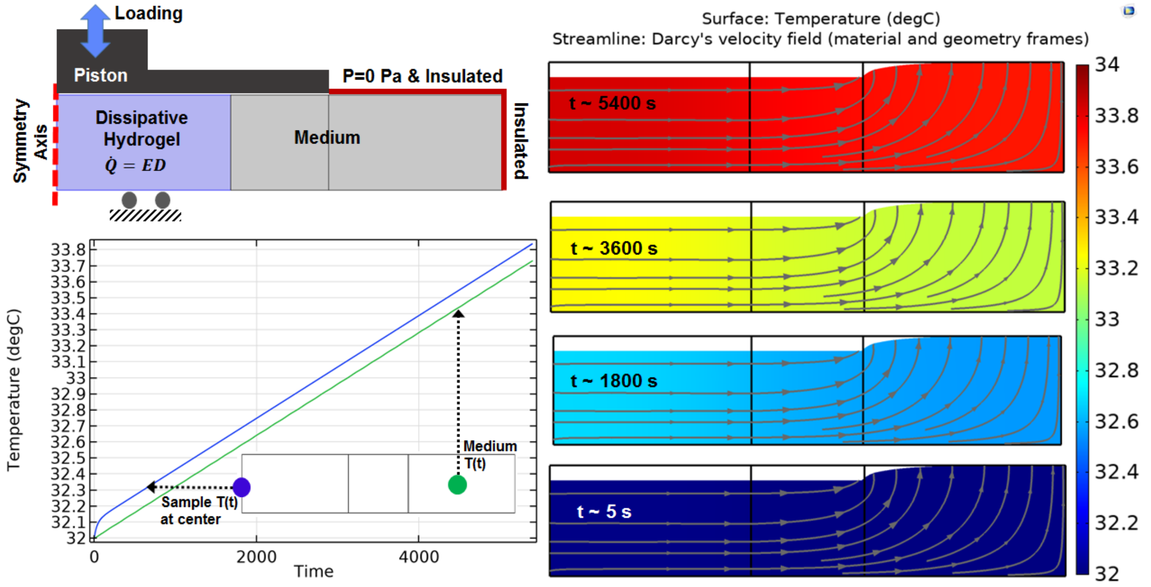


**Figure S3.** Heat transfer simulation in one culture well of the bioreactor by application of adiabatic buondary condition. The temperatue of the sample and medium was changed around 2°C over stimulation period showing that hydrogel dissipative capacity can cause self-heating in specific boundary condition. The color map illustrates temperature and the arrowed curves are velocity streamlines. Since the size of the sample is small compared to the medium, the generated heat by the sample can only increase temperature from 32°C to around 34°C.

**Analytical Heat Transfer Model**

By assuming a closed adiabatic system around dissipative hydrogel (excluding culture medium) and the hypothesis of local thermal equilibrium between fluid and solid phases, the temperature evolution of dissipative can be analytically calculated by a lumped heat transfer model as following:

$$(\rho C)_{eff} \frac{\partial T}{\partial t} = \dot{q}$$

$$(\rho C)_{eff} = \phi \rho_{fluid} C_{fluid} + (1 - \phi) \rho_{solid} C_{solid}$$

where $(\rho C)_{eff}$ is the effective volumetric heat capacity of the hydrogel and $\dot{q}$ represents the power of lost energy in each cycle of loading-unloading. This simplified model, predict about 10$^{o}$C temperature increase inside the hydrogel scaffold after one hour of cyclic loading.

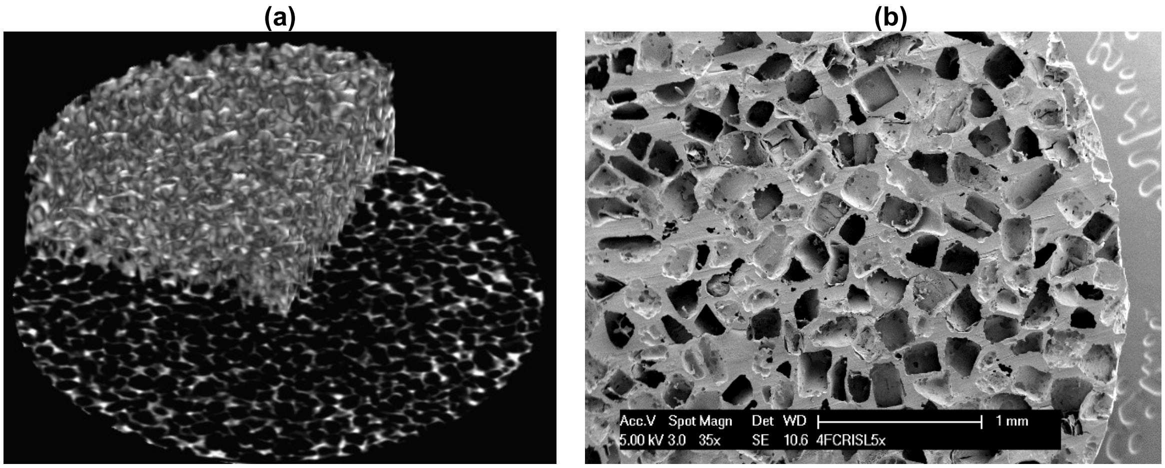


**Figure S4.** Morphological structure of the dissipative hydrogel. (a) Reconstructed micro CT scans. (b) scanning electron microscopy image of the hydrogel scaffold.

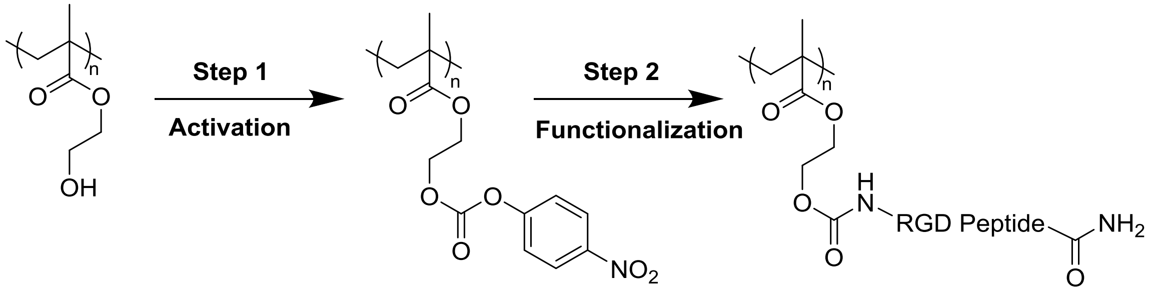


**Figure S5.** Applied two-step bio-conjugation process (hydroxyl group activation and RGD peptide grafting) for functionalization of pHEMA based hydrogels.

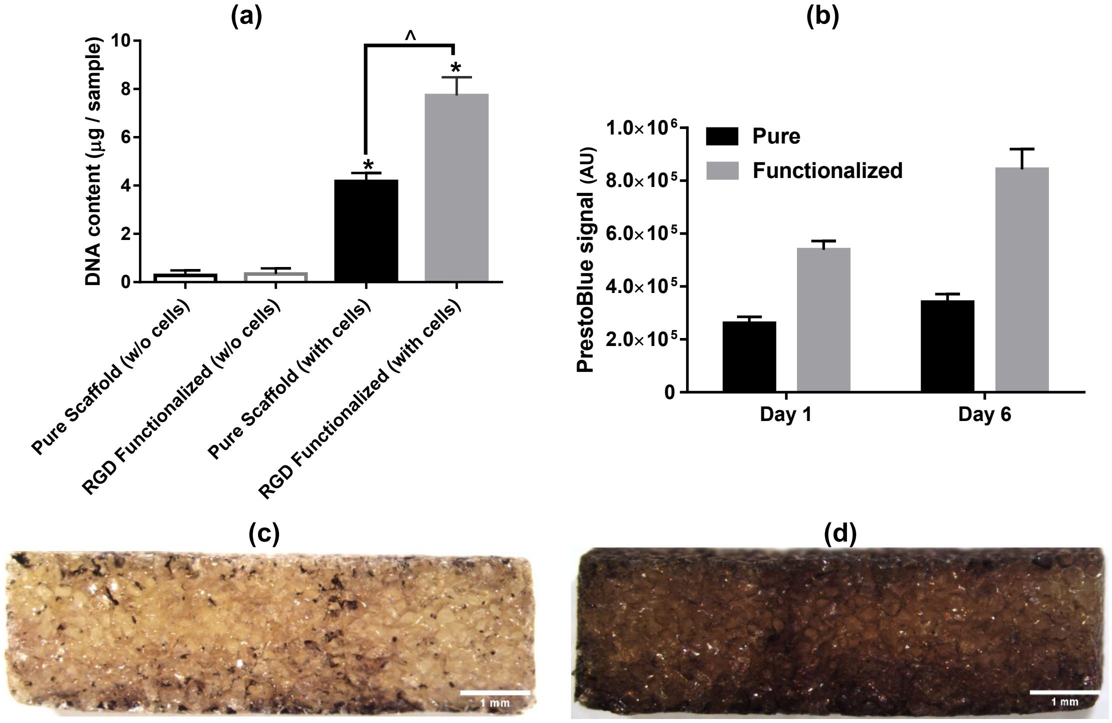


**Figure S6.** Cell attachment, proliferation and distribution inside pure (non-modified) and RGD functionalized hydrogels. (a) DNA content assay for cell-seeded hydrogels demonstrated significantly higher cell attachment in RGD functionalized samples compared to pure hydrogels ($p<0.05$, Student t-test, n=3). (b) Cells proliferation were significantly higher in RGD functionalized hydrogels over the static culture period compared to non-modified pure samples. (c,d) Cell distribution on central cross section of the hydrogel discs after 1 (top-right) and 10 days (bottom-right), respectively.The cells could penetrate inside porous hydrogels and grow overtime because of available binding sites by RGD peptieds.

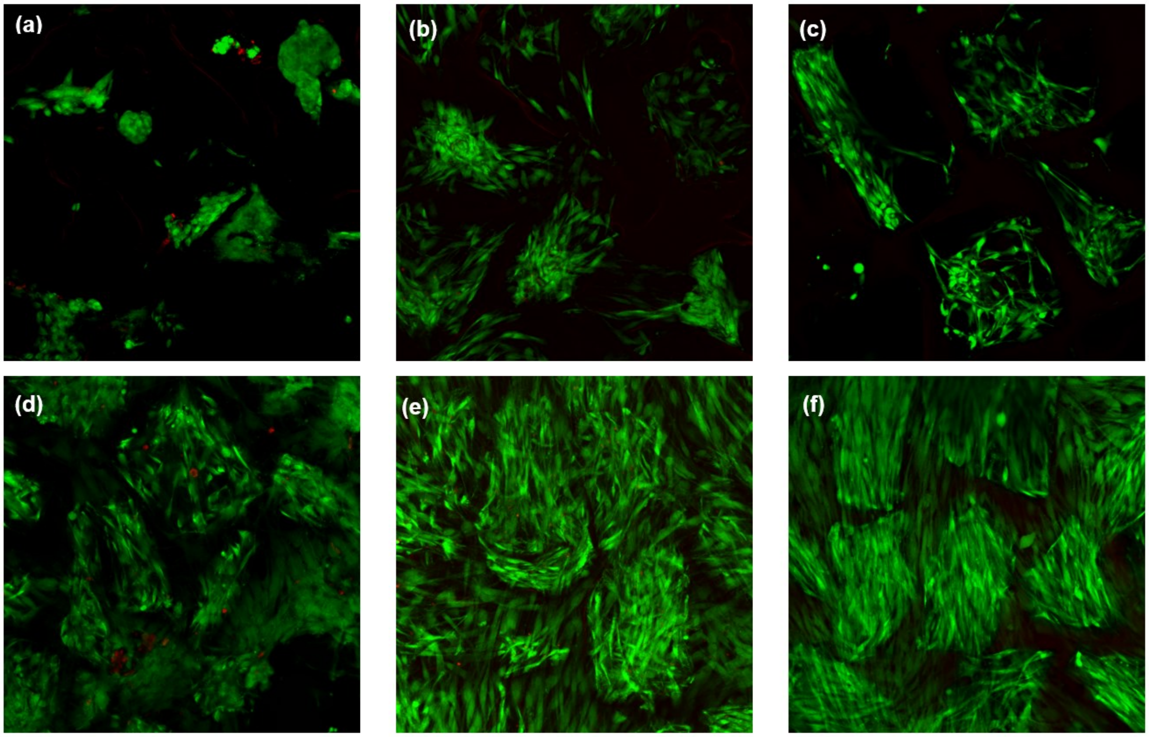


**Figure S7.** Cell viability and attachment before and after thermo-mechanical stimulation inside the bioreactor for non-modified pure (top) and RGD functionalized (bottom) hydrogels. (a , d) Cell viability for free-swelling samples at 32$^oC$ on day 2. (b , e) Cell viability for free-swelling samples at 32$^oC$ on day 10. (c , f) Cell viability after thermo-mechanical stimulation during 2 hours of 10 % cyclic compression at 1 Hz over 10% pre-strain along with temperature increase from 32 to 39$^oC$.

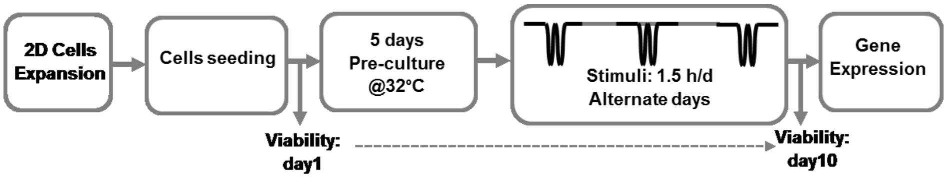


**Figure S8.** Schematic workflow of the performed *in vitro* thermo-mechanobiological experiment.

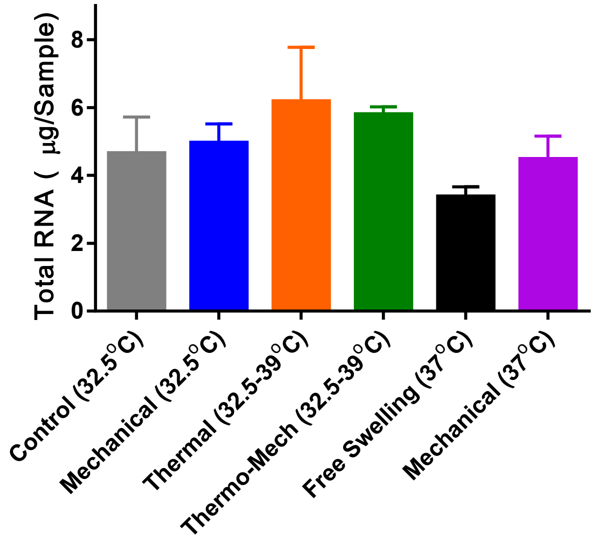


**Figure S9.** Total RNA as an indicator for cell metabolism was minimally varied by intermittent thermo-mechanical stimulation when culture baseline temperature was set at 32°C (knee temperature at rest). A decrease of total RNA for continuous incubation of cells at 37°C (core body temperature) was also observed.

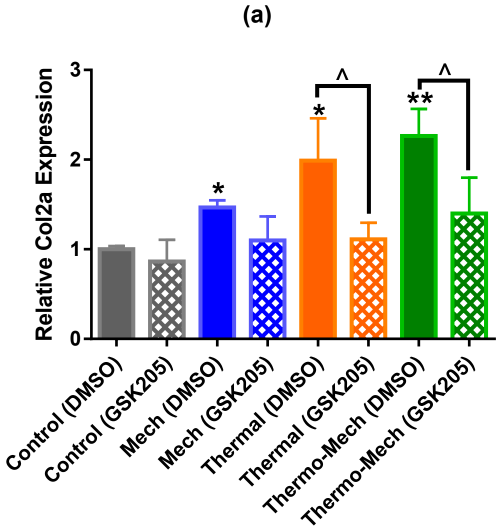


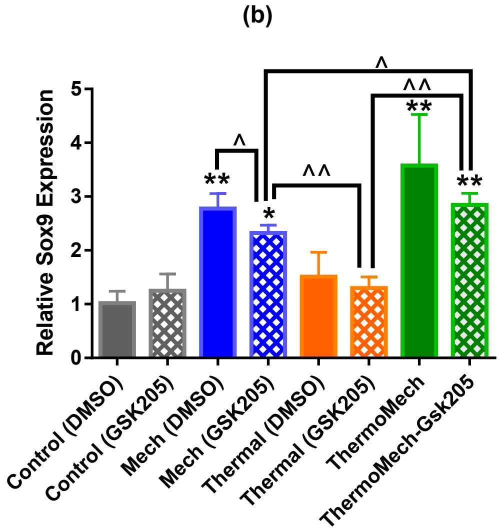


**Figure S10.** Gene expression results indicated that TRPV4 acts as signal integrator for thermal and mechanical stimuli. Inhibition of TRPV4 channels by GSK205, significantly reduced the expression of Col2a in Thermal group, and slightly in mechanical group. In parallel, TRPV4 channels disactivation, significantly reduced Sox9 expression in mechncially loaded samples and slightly in thermal group. This data confirms that TRPV4 channels are influential in transduction of thermal and mechanical cues and their contribution could be different deponding on the target indicator. In addition, significant upregulation of Sox9 in mechanically loaded samples with and without GSK205 could imply the contribution of other signal mediators such as integrins through cell-scaffold interaction with respect to incorporated RGD motifs.

**Table S1.** Primers data used for qRT-PCR.

| Primer Name | Sequence | Concent. | Efficiency | Temp |
|---|---|---|---|---|
| *RPL13-F* | TAAACAGGTACTGCTGGGCCG | 150 ng | 96 | 60 |
| *RPL13-R* | CTCGGGAAGGGTTGGTGTTC | | | |
| *Agc-F* | GGTACCAGTGCACAGAGGGGTT | 175 ng | 99 | 62 |
| *Agc-R* | TGCAGGTGATCTGAGGCTCCTC | | | |
| *Twist-F* | AGCAGGGCCGGAGACCTAGATGTCA | 250 ng | 95 | 60 |
| *Twist-R* | ACGGGCCTGTCTCGCTTTCTCT | | | |
| *Comp-F* | TGCTTCGGGAACTGCAGGAAAC | 250 ng | 101 | 60 |
| *Comp-R* | GCACGCGTCACACTCCATCACC | | | |
| *SOX9-F* | TGGAAACTTCAGTGGCGCGGA | 225 ng | 108 | 64 |
| *SOX9-R* | AGAGCAAAAGTGGGGGCGCTT | | | |
| *COL1A1-F* | CCTGCGTACAGAACGGCCTCA | 150 ng | 88 | 60 |
| *COL1A1 -R* | CGTCATCGCACAACACCTTGCC | | | |
| *Col2a- F* | GGAATTCGGTGTGGACATAGG | 175 ng | 96 | 60 |
| *Col2a- R* | ACTTGGGTCCTTTGGGTTTG | | | |

**Table S2.** Material properties used in the heat transfer model of bioreactor culture well.

| Material Properties | Value | Unit |
|---|---|---|
| *Equilibrium modulus (Eeq)* | 500 | kPa |
| *Poisson ratio(v)* | 0.23 | - |
| *Porosity (ϕ)* | 68 | % |
| *Permeability (k)* | $2.1e^{-14}$ | $m^2$ |
| *Dissipative Power of Hydrogel* | 9000 | $w/m^3$ |
| *Heat capacity of pHEMA (Csolid)* | 1308 | J/(kg.Kelvin) |
| *Conductivity of pHEMA (Ksolid)* | 0.25 | W/(m.Kelvin) |
| *Heat capacity of water, (Cfluid)* | 4200 | J/(kg.Kelvin) |
| *Conductivity of water, (Kfluid)* | 0.6 | W/(m.Kelvin) |
| *Biot–Willis Coefficient (α)* | 1 | - |
| *Relaxation modulus (G1)* | 200 | kPa |
| *Relaxation modulus (G2)* | 58 | kPa |
| *Relaxation modulus (G3)* | 215 | kPa |
| *Relaxation time ($\tau_1$)* | 0.42 | s |
| *Relaxation time ($\tau_2$)* | 5.82 | s |
| *Relaxation time ($\tau_3$)* | 1600 | s |

**Movie S1 (separate file).** TRPV4 channels activation in human chondro-progenitor cells cultured in 2D by using 10nM GSK01 agonist**.**

**Movie S2 (separate file).** TRPV4 channels inhibition in human chondro-progenitor cells in 2D by using 10 μM GSK205 antagonist**.**
